# A GYS2/p53 negative feedback loop restricts tumor growth in HBV-related hepatocellular carcinoma

**DOI:** 10.1101/373761

**Authors:** Shi-Lu Chen, Chris Zhiyi Zhang, Li-Li Liu, Shi-Xun Lu, Ying-Hua Pan, Chun-Hua Wang, Yang-Fan He, Cen-Shan Lin, Xia Yang, Dan Xie, Jing-Ping Yun

## Abstract

Hepatocarcinogenesis is attributed to the reprogramming of cellular metabolism as consequence of the alteration in metabolite-related gene regulation. Identifying the mechanism of aberrant metabolism is of great potential to provide novel targets for the treatment of hepatocellular carcinoma (HCC). Here, we demonstrated that glycogen synthase 2 (GYS2) restricted tumor growth in HBV-related HCC via a negative feedback loop with p53. Expression of GYS2 was significantly downregulated in HCC and correlated with decreased glycogen content and unfavorable patient outcomes. GYS2 overexpression suppressed, whereas GYS2 knockdown facilitated cell proliferation *in vitro* and tumor growth *in vivo* via modulating p53 expression. GYS2 competitively bound to MDM2 to prevent p53 from MDM2-mediated ubiquitination and degradation. Furthermore, GYS2 enhanced the p300-induced acetylation of p53 at K373/382, which in turn inhibited the transcription of GYS2 in the support of HBx/HDAC1 complex. In summary, our findings suggest that GYS2 serves as a prognostic factor and functions as a tumor suppressor in HCC. The newly identified HBx/GYS2/p53 axis is responsible for the deregulation of glycogen metabolism and represents a promising therapeutic target for the clinical management of HCC.

**Synopsis:** This study elucidate the role of GYS2 in glycogen metabolism and the progression of HCC. The newly identified HBx/GYS2/p53 axis is responsible for the deregulation of glycogen metabolism and represents a promising therapeutic target for the clinical management of HCC.

1. Decrease of GYS2 was significantly correlated with decreased glycogen content and unfavorable patient outcomes in a large cohort containing 768 patients with HCC.
2. GYS2 overexpression suppressed, whereas GYS2 knockdown facilitated cell proliferation in vitro and tumor growth in vivo via modulating p53 signaling pathway.
3. GYS2 competitively bound to MDM2 to prevent p53 from MDM2-mediated ubiquitination and degradation.
4. GYS2 enhanced the p300-induced acetylation of p53 at Lys373/382, which in turn inhibited the transcription of GYS2 in the support of HBx/HDAC1 complex.

## Introduction

Hepatocellular carcinoma (HCC) ranks the fifth most prevalent malignancies and the second leading cause of cancer-related death worldwide (Ferlay et al, 2015). Epidemiological surveys indicate that more than half of HCC occurs on a background of hepatitis B virus (HBV)-related immflammation and metabolomic alterations (Cvitanovic et al, 2017; Tang et al, 2018). Emerging evidence has strongly suggested that aberrant glucose metabolism is a highlighted hallmark of cancers (Bian et al, 2017; Boroughs & DeBerardinis, 2015), yet the abnormalities in glycogen regulation has rarely been studied (Morris, 2018; Upadhyay et al, 2013).

Glycogen is a branched polymer of glucose that is stored in muscle and liver to serve as an energy supply in times of need (Roach et al, 2012; Stalmans et al, 1987). Aberrant glycogen content has been observed in 58 kinds of cultured human tumor cell lines: increased in cells originated from low glycogen tissues such as breast, ovary, kidney and lung, but decreased in HCC cell line SK-Hep-1 and choriocarcinoma JEG-3 (Lea et al, 1972; Rousset et al, 1981). An inverse correlation between glycogen concentration and tumor growth was observed (Rousset et al, 1981). For example, loss of glycogen debranching enzyme AGL led to the reduction of glycogen and enhancement of bladder cancer growth independent of its enzymatic activity (Guin et al, 2014). In glioblastoma, breast and colon cancer cells, hypoxia induced glycogen accumulation, premature senescence and tumor growth suppression in a p53-dependent manner (Favaro et al, 2012). However, the alteration of glycogen metabolism in HCC and the underlying molecular mechanism remain largely unknown.

Glycogen synthase (GS) is a key enzyme in glycogen synthesis. There are two isoforms of GS, glycogen synthase 1 (GYS1) and glycogen synthase 2 (GYS2) in human tissues. It is well established that GYS1 is generally expressed in muscle, heart and kidney, while GYS2 expression appears to be primarily restricted to the liver (Nuttall et al, 1994; Roach et al, 2012). As the rate-limiting enzyme of glycogen biosynthesis, GS catalyzes the addition of UDP-glucose onto existing glucose molecules to elongate linear glucose polymer chain. Previous studies indicated that GYS2 deficiency caused glycogen storage disease type 0 (GSD-0) with the symptom of impaired glucose tolerance (Orho et al, 1998; Szymanska et al, 2015). GYS2 was phosphorylated and suppressed by glycogen synthase kinase 3β (GSK3β), allosterically activated by glucose-6-phosphate (G6P) (von Wilamowitz-Moellendorff et al, 2013; Wang et al, 2012) and transcriptionally regulated by CLOCK gene in the circadian rhythms of hepatic glycogen synthesis (Doi et al, 2010). The role of GS in the progression of cancer cells is rarely studied. GYS1 was required for glycogen flux and myeloid leukemia cell growth via activating AMPK pathway (Bhanot et al, 2015). The PI3K/AKT-mediated phosphorylation of GYS2 was induced by mulberry anthocyanin extract in HepG2 cells (Yan et al, 2016). However, the clinical significance of GYS2 and its bio-function in glycogen regulation in human cancers are still unclear.

In the present study, we aimed to elucidate the role of GYS2 in glycogen metabolism and the progression of HCC. Our data showed for the first time that decreased expression of GYS2 resulted in the reduction of glycogen and indicated unfavorable clinical outcomes. *In vitro* and *in vivo* experiments demonstrated that GYS2 restricted tumor growth in HBV-related HCC via a negative feedback loop with p53. Our findings supplement the understanding of HCC glycogen metabolism and provide potential prognostic and therapeutic targets for HCC treatment.

## Results

### Glycogen content is decreased in HCC

Using Periodic acid-Schiff (PAS) staining, we noticed a marked decrease of glycogen in HCC tissues, compared with the nontumorous tissues (Fig. 1A). Quantitative colorimetric results confirmed the decreased amount of glycogen content in 24 HCC tissues (Fig. 1B). In a large cohort consisting of 768 HCC patients, 60.68% (466/768) of the cases showed weaker PAS staining in tumor tissues, while HCC tissues contained more glycogen in 20.18% (155/768) of HCC cases (Fig. 1C). We next determine the clinical significance of glycogen content in HCC. Patients were divided into high and low PAS groups according to the positive staining proportion. Kaplan-Meier analyses revealed that low PAS staining was correlated with unfavorable overall survival (Fig. 1D). Significant association between low PAS staining and HBV infection, larger tumor size, advanced TNM stage and poorer tumor differentiation was identified (Supplementary Table 2). Multivariate cox regression model further indicated glycogen content as an independent prognostic factor of overall survival in HCC (Supplementary Table 3). These data showed that glycogen synthesis was inhibited in HCC.

**Figure 1.**
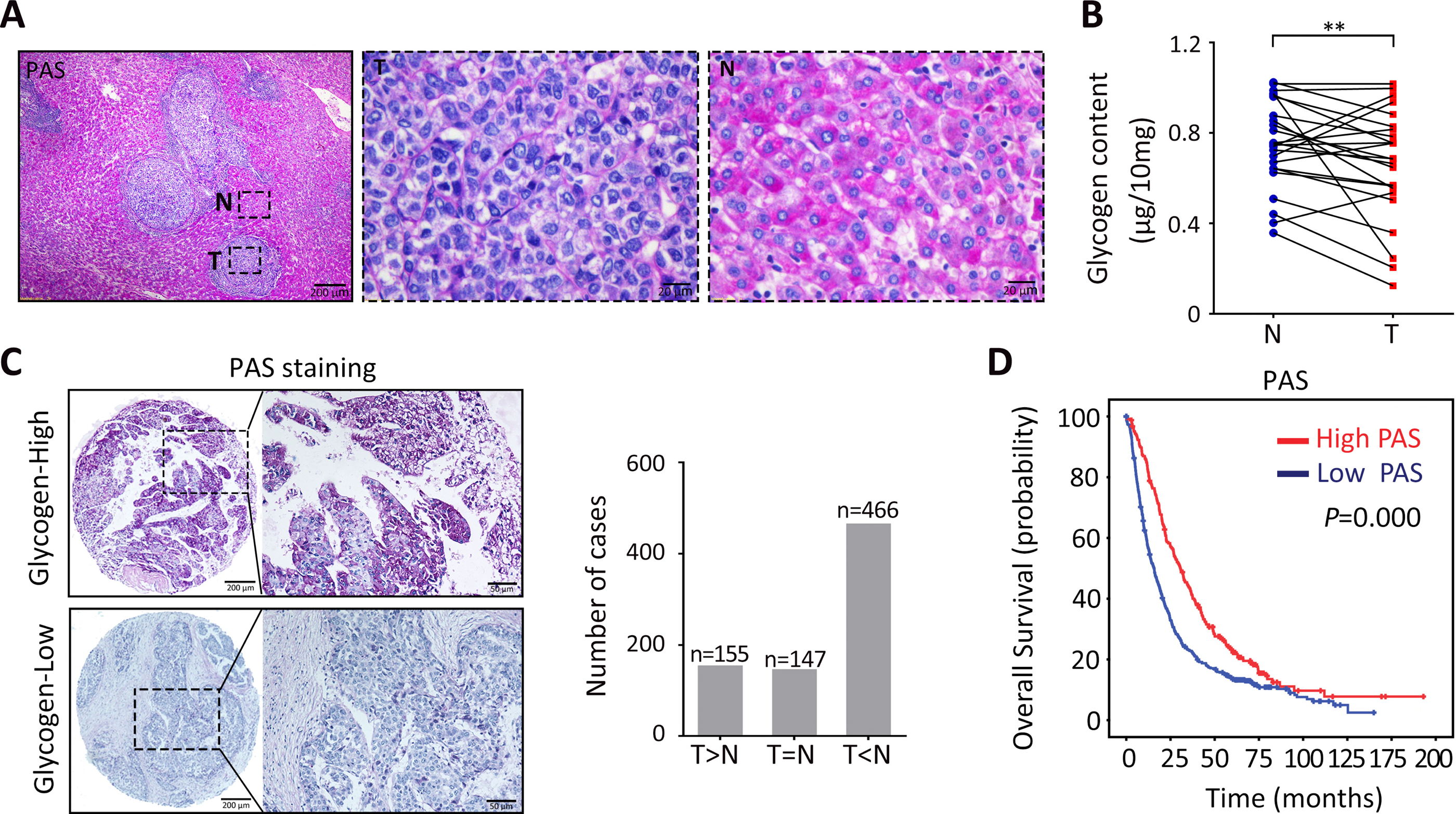
Glycogen is decreased in HCC and associated with disease progression. **A.** PAS staining was used to detect the glycogen content in formalin-fixed liver tissues. Representative images of Tumor (T) and non-tumor (N) in one HCC specimen were presented. **B.** Glycogen levels were quantified by quantitive colorimetric method in 24 HCC and corresponding adjacent liver tissues. Fresh tissue weight was used to normalize as ratio. **C.** The glycogen content in a HCC cohort of 768 patients were determined by PAS staining. Representative images of tumor tissues in two groups were shown. Tumor and corresponding non-tumor glycogen content were compared and the number of cases in each group were indicated. **D.** The TMA cohort was separated into high and low glycogen groups according to PAS staining. The correlation of PAS staining and overall survival was determined by Kaplan-Meier analysis. Statistical data were represented as mean± SD. ^**^*P*<0.01.

### GYS2 is decreased in HCC and correlated with poor patient outcome

As shown by the transcriptional profiling in 8 paired HCC tissues (GSE104310), genes involved in glycogen synthesis and metabolism were deregulated. Specially, GYS2 was mostly downregulated in tumor tissues (Fig. 2A), which was verified in a large cohort of HCC patients. GYS2 mRNA and protein levels in HCC cell lines were lower than immortalized hepatic cell L-02 (Supplementary Fig. 1a). In 48 pairs of HCC fresh tissues, GYS2 mRNA was reduced compared to matched non-tumor tissues (Fig. 2B). Oncomine data supported the decrease of GYS2 mRNA (Supplementary Fig. 1b). Consistently, GYS2 protein level was significantly lower in HCC specimens, and positively correlated with glycogen content (Fig. 2C-D). Examination of GYS2 expression in 763 HCC cases showed that GYS2 was remarkably downregulated in tumor tissues (Fig. 2E). Patients were divided into high and low GYS2 groups according to the median IHC score (4.0). Low GYS2 expression, identified in 60.6% (462/763) of HCC tissues, was frequently observed in the cases with weak PAS staining (Fig. 2F). Significant association was found between low GYS2 expression and higher serum AFP level, larger tumor size and HBV infection (Supplementary Table 4). Kaplan-Meier analyses revealed that patients with GYS2 deficiency was accompanied with unfavorable overall survival in both SYSUCC and TCGA cohorts (Fig. 2G). Multivariate cox regression model further indicated glycogen content as an independent prognostic factor of overall survival in HCC (Supplementary Table 5). This was further validated by stratified survival analyses (Supplementary Fig. S2). The expression of GYS2 was next determined in the HCC metastatic nodules in portal vein (Supplementary Fig. 3A). No significant correlation was found between GYS2 expression and the disease-free survival (Supplementary Fig. 3B).

**Figure 2.**
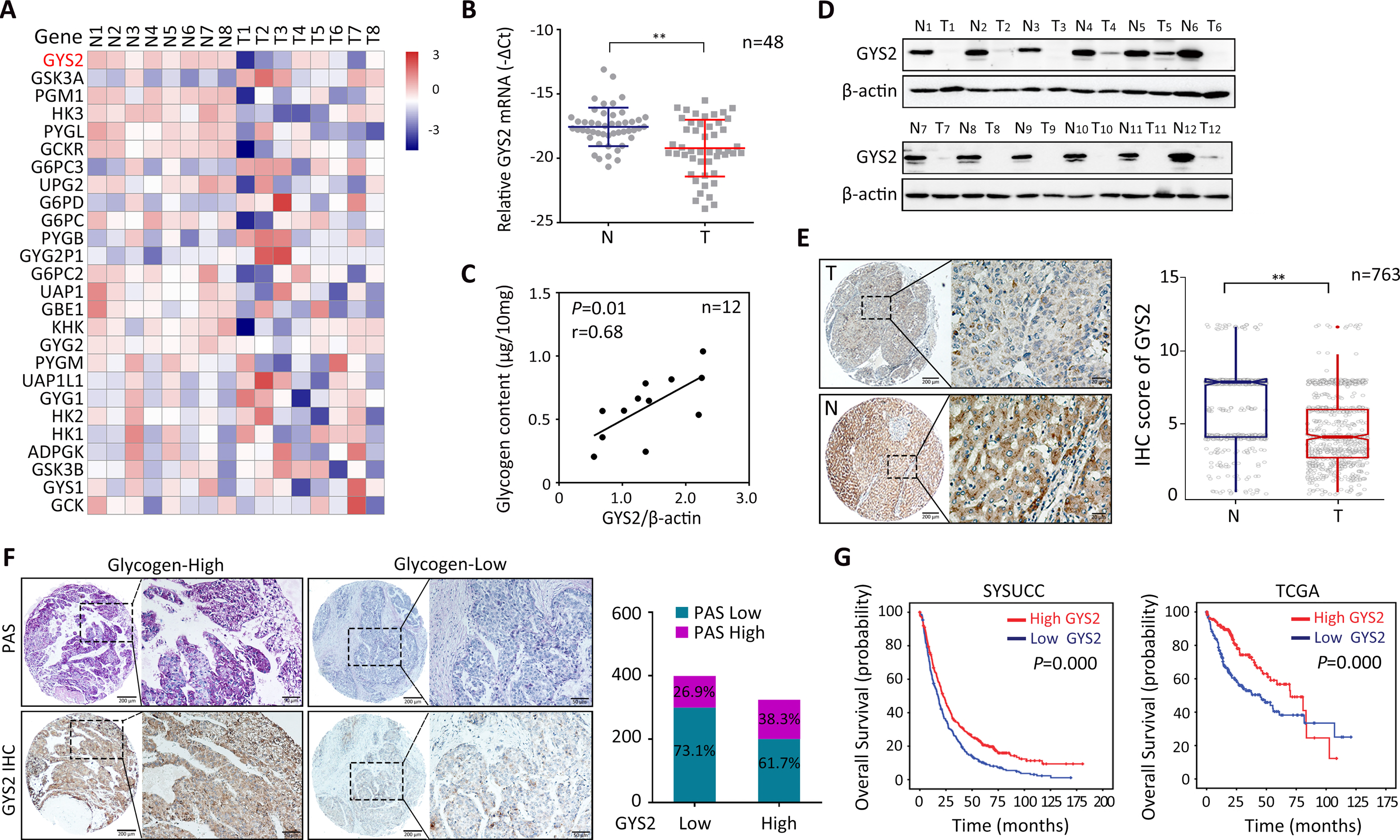
GYS2 is decreased in HCC and correlated with poor patient outcome. **A.** RNA-seq in 8 paired tissues were conducted to identify the variation of genes in HCC. Heat map depicted the mRNA level of genes involved in glycogen synthesis and metabolism. Blue denotes lower expression and red for elevated expression. **B.** The mRNA expression of GYS2 in 48 pairs of HCC and corresponding adjacent liver tissues were determined. 18S RNA was used to normalize the fold change. **C.** Expression profile of GYS2 protein in 12 paired HCC and adjacent non-tumor tissues was detected by western blot. β-actin was used to indicate the amount of loading proteins. **D.** The glycogen content was determined by quantative colorimetric method in 12 pair of HCC tissues and the correlation with GYS2 protein were analyzed. **E.** The expression of GYS2 was determined by TMA-based IHC staining. The representative images of tumor (T) and non-tumor (N) with the score of each cases were shown. **F.** The expression of GYS2 was presented by IHC staining and glycogen content were detected by PAS staining. The correlation between GYS2 and PAS were assessed and PAS proportion were shown as ratio in high and low GYS2 groups. **G.** The correlation between GYS2 expression and overall survival were determined in SYSUCC cohort and TCGA cohort by Kaplan-Meier analysis. Statistical data were represented as mean± SD. ^**^*P*<0.01.

### Loss of GYS2 promotes HCC proliferation *in vitro* and *in vivo*

To explore the biological function of GYS2 in HCC, GYS2 was either knocked down by siRNA in HepG2 and QGY-7703 cells or overexpressed in Bel-7402 and Bel-7404 cells (Fig. 3A). PAS staining and quantitative assays showed that depletion of GYS2 decreased, whereas overexpression of GYS2 increased the glycogen amount in HCC cells (Supplementary Fig. 4A-B). Cell viability was markedly increased in GYS2-silenced cells but decreased in GYS2-expressing cells (Fig. 3B). EdU-positive cells were noticeably induced by GYS2 knockdown, but reduced by GYS2 overexpression (Fig. 3C). Cells were arrested at S/G2/M phase upon the silence of GYS2. Overexpression of GYS2 resulted in more cells at G0/G1 phase compared to control group (Fig. 3D). Furthermore, GYS2-depleted cells formed more colonies. In contrast, ectopic expression of GYS2 weakened the HCC cell proliferation (Fig. 3E). However, the alteration of GYS2 expression led to no change of cell apoptosis and cell migration (Supplementary Fig. 5A-B). To validate these effects *in vivo*, we established a xenograft model by subcutaneously injecting cells into of nude mice. Tumors bearing GYS2-silencing cells grew faster, contained less glycogen and expressed higher Ki-67, compared to the control group. Conversely, mice injected with GYS2-expressing cells carried smaller tumors that were with higher glycogen content and lower Ki-67 expression (Fig. 3F & Supplementary Fig. 6). Collectively, these findings indicate that lack of GYS2 greatly contributes to HCC cell growth.

**Figure 3.**
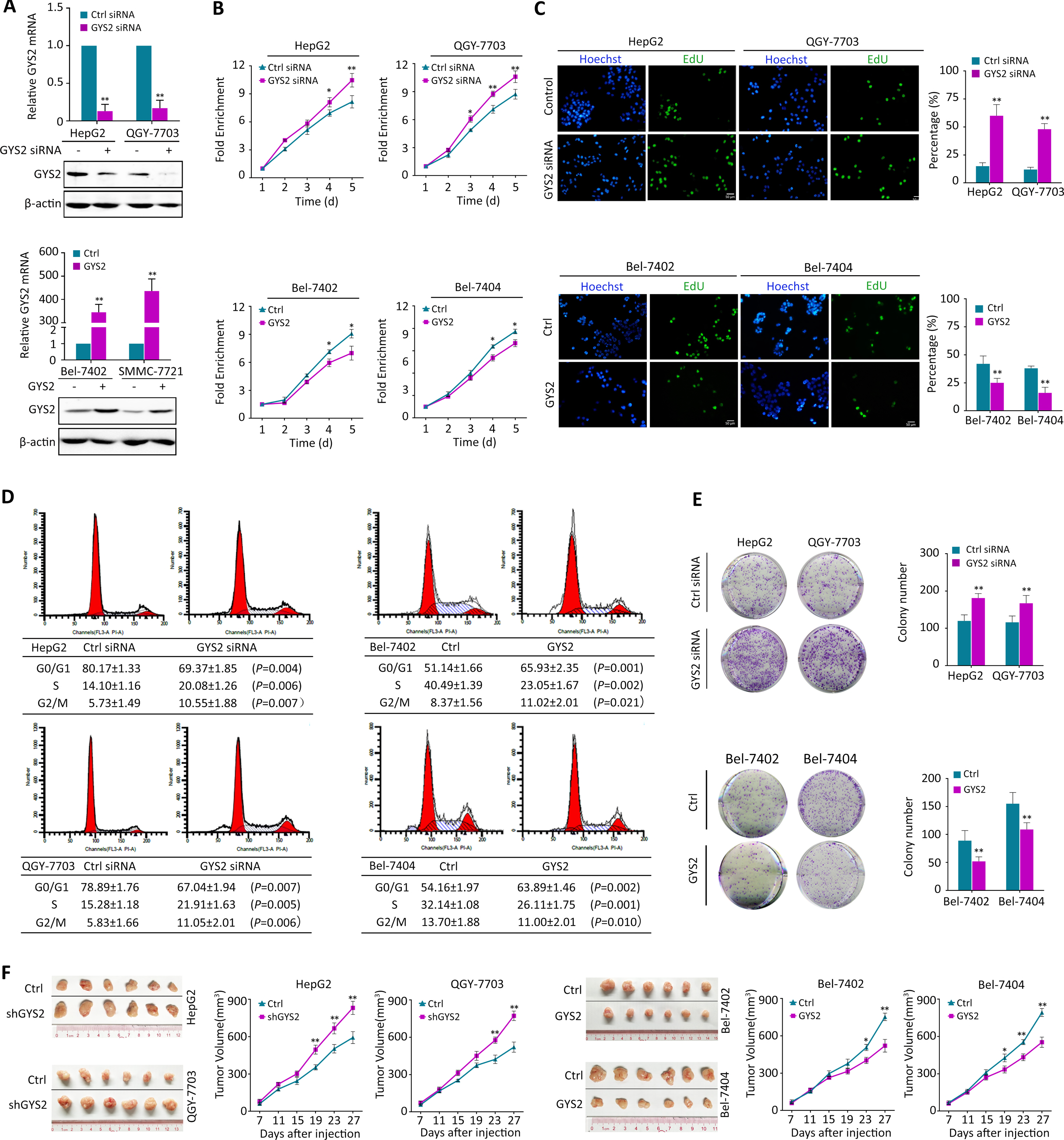
Loss of GYS2 promotes HCC proliferation *in vitro* and *in vivo.* **A.** GYS2 was silenced using siRNA in HepG2 and QGY-7703 cells or exogenous expressed by transfecting pcDNA3.1-GYS2 plasmids to Bel-7402 and Bel-7404 cells. The mRNA and protein levels of GYS2 were determined by qRT-PCR and Western blot. **B.** Cell proliferation rates in GYS2-depletion (HepG2 and QGY-7703) or overexpressed (Bel-7402 and Bel-7404) cells were detected by MTT assay in five consecutive days. Relative absorbance was measured at OD490. Fold changes in each day were normalized to the absorbance record in day 1. **C.** EdU assay showed the replication of DNA in cells induced by GYS2 knockdown or overexpression. Green denotes duplicated cells and blue denotes cell nucleus. **D.** GYS2 was silenced or overexpressed in indicated cells. Flow cytometry assays determined the percentage of cells in cell cycle. **E.** Colony formation assays were used to determine the effect of GYS2 on cell growth. 1000 cells in each groups were seeded into 6-well plate and 14 days later, the number of colony were counted using Image J software.**F.** Xenograft mice experiment was carried out to determine the tumor growth *in vivo*. Mice were sacrificed 27 days after injections of HCC cells stably silencing or overexpressing GYS2. The images of tumors in each groups were shown and tumor volume were calculated. Statistical data were represented as mean± SD. ^*^*P*<0.05, ^**^*P*<0.01.

### GYS2 activates p53 signaling pathway

To unveil the underlying mechanism of GYS2-mediated HCC proliferation, we conducted transcriptional profiling by RNA-sequencing in HepG2 and QGY-7703 cells. Results showed that p53 pathway was significantly suppressed in both cell lines with GYS2 depletion (Fig. 4A). Knockdown of GYS2 reduced, whereas overexpression of GYS2 induced the p53 protein level, resulting in the modulation of p53-targeted genes, such as p21, 14-3-3σ and Cyclin D1 (Fig. 4B). Because GYS2 had no effect on the transcriptional regulation of p53 (Supplementary Fig. 7A), we next examined the protein interaction between the two proteins. Co-IP data revealed that GYS2 was detectable in p53 antibody-mediated precipitate in the cytoplasm (Fig. 4C). GST-pull down assay verified the direct binding of GYS2 and p53 (Fig. 4D). The co-localization between GYS2 and p53 was primarily observed in cytoplasm by confocal assays (Fig. 4E). We further determined the interacting domains. Results demonstrated that the regions 1-101 aa of p53 and 1-500 aa of GYS2 were required for the GYS2-p53 binding (Fig. 4F&G and Supplementary Fig.7B).

**Figure 4.**
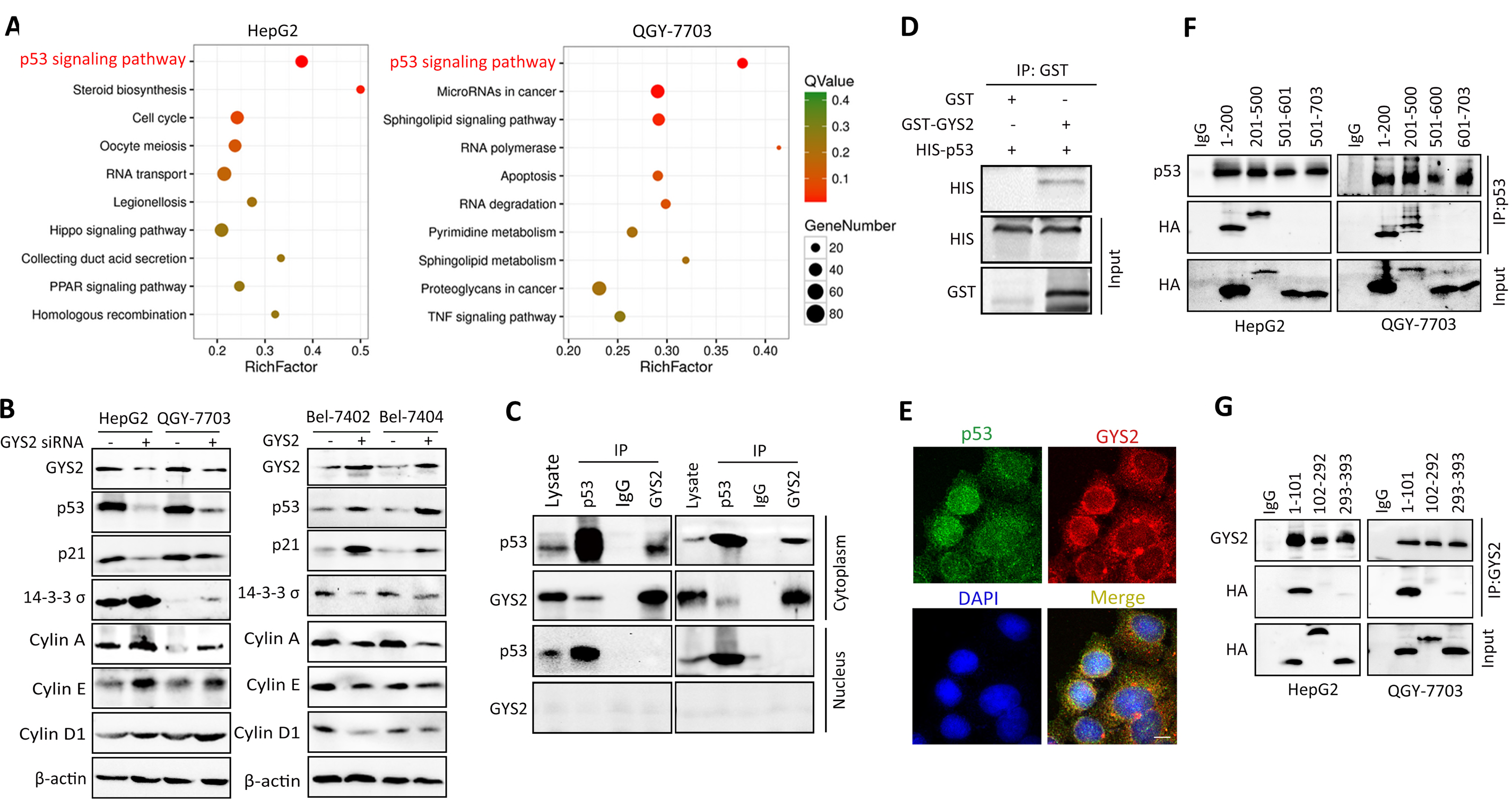
GYS2 activate p53 signaling pathway. **A.** RNA-seq were carried out to identify the downstream target of GYS2. KEGG enrichment analysis showed the transcriptional profiling of enriched pathways in GYS2-depleted HepG2 and QGY-7703 cells. **B** GYS2 was silenced using siRNA in HepG2 and QGY-7703 or overexpressed in Bel-7402 and Bel-7404. Subsequently, the protein expression of p53 and its downstream targets were determined by western blot. **C.** Co-IP assays were performed to determine the interaction of GYS2 and p53 in cytoplasm and nucleus. **D.** GST-pull down assays were carried out to determine the direct interaction between GST-tagged GYS2 and His-tagged p53. **E.** IF staining indicating the co-localization of p53 (green) and GYS2 (red) together with DAPI (blue) in HepG2 cells. **F.G.** The critical sites for the interaction between GYS2 and p53 were measured in HepG2 and QGY-7703 cells overexpressed with truncated GYS2 plasmids (F) or p53 plasmids (G).

To validate if GYS2 exerted anti-HCC activities via p53, we carried out rescue experiments. Restoring the expression of p53 partly attenuated the cell growth promoted by GYS2-depletion in HepG2 and QGY-7703. By contrast, p53 knockdown rescue the inhibitory effect of GYS2-expression on cell proliferation (Supplementary Fig. 8A-B). These data suggest that GYS2 functions as a tumor suppressor via interaction with p53 in HCC cells.

### GYS2 stabilizes p53 via competitive interaction with MDM2

We next explored the mechanism via which GYS2 upregulated p53. Using cyclohexamide (CHX, a translation inhibitor), we found that GYS2 overexpression markedly prolonged the half-life of p53 protein in HCC cells (Fig. 5A). In cells with GYS2 depletion, p53 was degraded much faster, compared with the control cells (Supplementary Fig. 9A). The ubiquitin-mediated proteasomal degradation of p53 was enhanced by GYS2 silence, but attenuated by ectopic GYS2 expression (Fig. 5b and Supplementary Fig. 9B). The GYS2 siRNA-induced p53 reduction was abolished by proteasome inhibitor MG132 (Fig. 5C).

**Figure 5.**
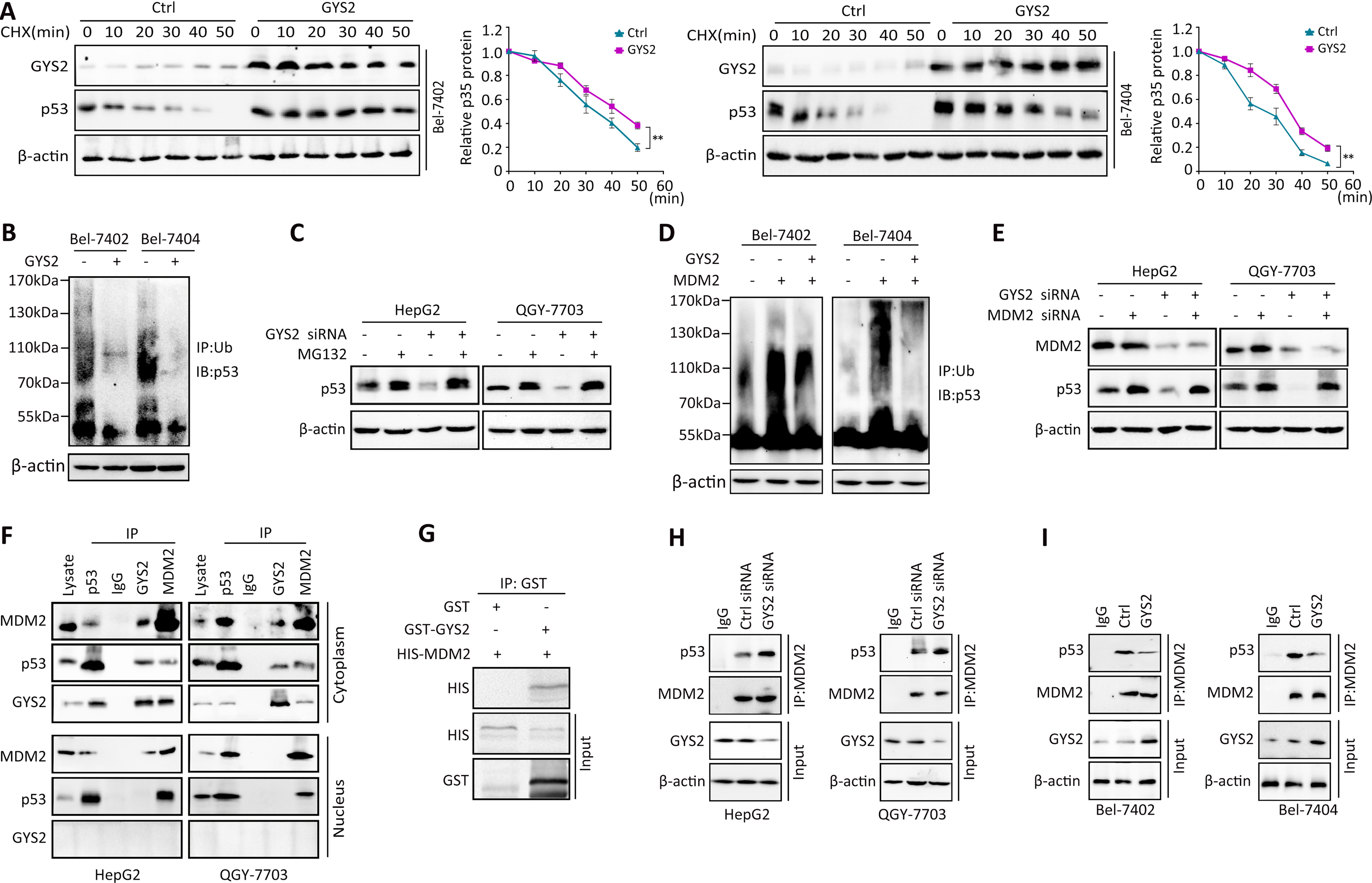
GYS2 stabilizes p53 via competitive interaction with MDM2. **A.** The half-life of p53 protein was detected in GYS2 overexpressed cells by added with CHX (20 μg/ml) for different times. The amount of p53 protein were quantitated and calculated by Image J software. **B.** Cells were pre-incubated with MG-132 (20 μM) for 12 h. Ub was immunoprecipitated (IP) and immunoblotted (IB) by anti-p53. The ubiquitination of p53 protein was detected after GYS2 overexpression. **C.** Cells were transfected with GYS2 siRNA and pre-incubated with MG-132 (20 μM) for 12 h. Cell lysate were immunoprecipitated by anti-Ub and immunoblotted by anti-p53. **D.** With the induction of MDM2, the effect of GYS2 on p53 ubiquitination were detected again as described by Figure 5 b. **E.** Cells were treated with MDM2 siRNA followed by GYS2 knockdown. p53 protein expression were detected by western blot. **F.** The interaction of GYS2, MDM2 and p53 was measured in HepG2 and QGY-7703 cells. **G.** GST-pull down detected the direct interaction between GST-tagged GYS2 and His-tagged MDM2. **H.I.** The interaction amount of p53 and MDM2 was measured after GYS2 knockdown (H) or overexpression (I). β-actin was used to indicate the amount of loading proteins. Statistical data were represented as mean± SD. ^**^*P*<0.01.

The E3 ubiquitin ligase MDM2 plays pivotal role in the p53 ubiquitination, we next determined whether MDM2 was involved in the regulation p53 by GYS2. MDM2-mediated p53 ubiquitination was partially attenuated by GYS2, and MDM2 siRNA-induced suppression of p53 ubiquitination was blocked by GYS2 siRNA (Fig. 5d and Supplementary Fig. 9C). Inhibition of MDM2 by siRNA or Nutlin-3 noticeably suppressed the p53 downregulation by GYS2 siRNA (Fig. 5E & Supplementary Fig. 9D). Since MDM2 bound to the 1–42 aa of p53 which was also the binding site for GYS2, we next determined the interaction among these three proteins. As shown by co-IP results, GYS2, p53 and MDM2 formed a protein complex through binding to each other (Fig. 5F). GST-pull-down assay also confirmed the direct binding of GYS2 and MDM2 (Fig. 5G). Notably, the interaction between MDM2 and p53 was enhanced by the knockdown of GYS2, but decreased by the overexpression of GYS2 in HCC cells (Fig. 5H-I). Taken together, these data indicate that GYS2 stabilizes p53 protein by competitively binding to MDM2 to inhibit the ubiquitination of p53.

### p53 represses GYS2 via a p300-dependent negative feedback loop

Previous studies demonstrated that p53 participated in metabolic regulation. We found that p53 was capable of transcriptionally regulating genes involved in glycogen synthesis, such as PYGL, GBE1, G6PC and GSK3β (Supplementary Fig. 10A). PAS staining and quantitative glycogen assays showed that overexpression of p53 reduced, whereas silencing of p53 induced the cytosolic glycogen content in HepG2 and QGY-7703 cells (Fig. 6A & Supplementary Fig. S12B). However, this effect was abolished in cells with GYS2 depletion (Supplementary Fig. 10C). These data prompted us to disclose the role of GYS2 in p53-mediated aberrant glycogen metabolism. Ectopic expression of p53 downregulated, while knockdown of p53 upregulated GYS2 at both mRNA and protein levels (Fig. 6B). Dual-luciferase assay showed that p53 transfection decreased the activity of GYS2 promoter (Fig. 6C). ChIP assays confirmed the binding of p53 to the GYS2 promoter (Fig. 6D).

**Figure 6.**
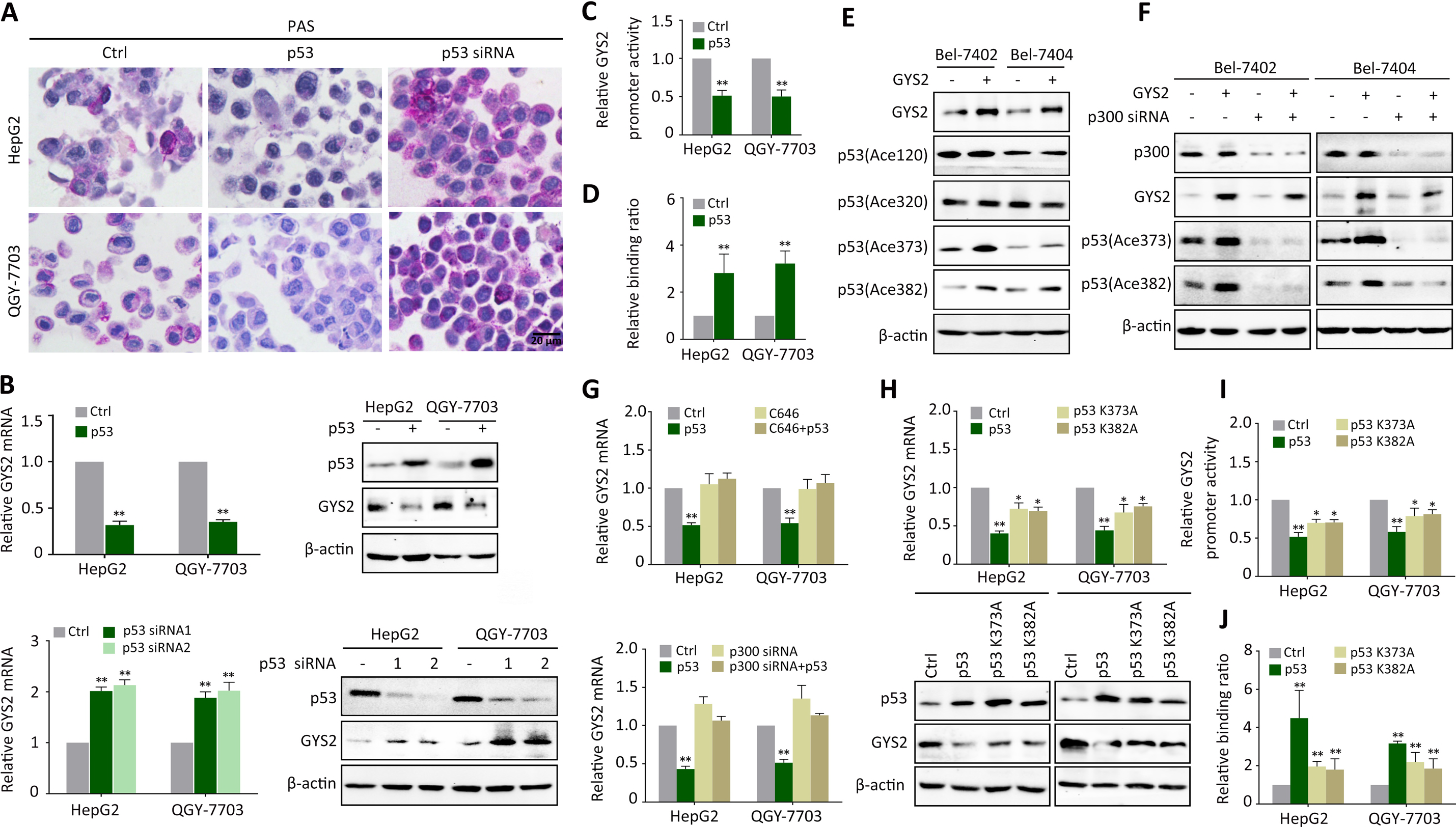
p53 represses GYS2 via a p300-dependent negative feedback loop. **A.** PAS staining showed the glycogen content in cells with p53 silenced or overexpression. **B.** The mRNA and protein expression of GYS2 were determined by qRT-PCR and western blot with the overexpression or silence of p53. **C. D.** The promoter activity of GYS2 was determined by dual-luciferase assay (C) and the binding ratio of p53 on GYS2 promoter was measured by ChIP (D). **E.** The expression of four p53 acetylated lysine sites were detected after GYS2 transfection. **F.** The expression of acetylated p53 in K373/382 were detected with the depletion of p300 using siRNA and overexpression of GYS2. **G.** The mRNA levels of GYS2 were detected while adding C646 (10 μM) or p300 depletion. **H.** The expression of GYS2 mRNA and protein was measured by qRT-PCR and western blot with the overexpression of p53 mutation in K373/382. **I.J.** Dual-luciferase assay (I) and ChIP (J) were carried out again to determine the effect of p53 K373/382 mutation on GYS2 promoter regulation. Statistical data were represented as mean± SD. ^*^*P*<0.05, ^**^*P*<0.01.

It is well-known that post-transcriptional modification of p53 is critically required for its transcriptional function. For example, p300-mediated acetylation of p53 enhances its DNA-binding ability to transactivate targets. We found that GYS2 overexpression increased p53 acetylation at lysine 373/382 (Fig. 6E), which was blocked by the knockdown of p300 (Fig. 6F). Given that the transcription activity of p53 is usually enhanced acetylation, we next determined the role of acetylated p53 in the suppression of GYS2 transcript. Treatment of cells with C646, a specific p300 inhibitor, or p300 siRNA, abolished the p53-mediated GYS2 decrease (Fig. 6G). We further constructed p53^K373A^ and p53^K382A^ mutations which cannot be acetylated by p300. Transfections of these two p53 mutants led to less reduction of GYS2 mRNA and protein (Fig. 6H). The inhibitory effect and the binding of p53 on GYS2 promoter was attenuated in cells with p53^K373A^ or p53^K382A^ expression (Fig. 6I-J). These results suggest that p53, in a negative feed-back loop, transcriptionally represses GYS2 via p300-mediated acetylation.

### HBx-HDAC1 complex facilitates the p53-mediated suppression of GYS2

Previous studies reported the involvement of HBV in liver glycogen regulation with conflicting results (Lu et al, 2013; Toshkov et al, 1994). But the role of HBx, the key oncogenic protein encoded by HBV, in glycogen metabolism remains unknown. We found that HBx overexpression decreased, while knockdown of HBx recovered the glycogen content in HCC cells (Fig. 7A). These effects were significantly rescued by GYS2 siRNA (Supplementary Fig. 11A). Our and Oncomine data suggested that GYS2 was further decreased in HBV-positive HCC patients (Supplementary Fig. 11B). Next, we intended to determine whether HBx played a role in the regulation of GYS2 by p53 in HCC cells. Reverse correlation between GYS2 and HBx was detected by TMA-based IHC (Fig. 7C). Low expression of GYS2 and PAS staining was more frequently found in HBx-positive cases (Fig. 7B and Supplementary Fig 12A). Silence of HBx in HBx-expressing HepG2.2.15 and HepG2-HBx cells resulted in induction of GYS2. On the other hand, HBx introduction in HBx-negative HepG2 and QGY-7703 cells downregulated GYS2 (Fig. 7C). Deletion fragments of HBx were constructed to identify the domain by which HBx suppressed GYS2 expression (Supplementary Fig. 12B). Results of qRT-PCR and western blot showed that overexpression of HBx C-terminal, but not N-terminal fragment, was capable of decreasing GYS2. Further data demonstrated that ΔHBx134 (C-terminal 20 aa truncated), but not ΔHBx120 (C-terminal 34 aa truncated), retained the suppressive effect of HBx on GYS2 (Supplementary Fig. 12C-D). These data indicate that the HBx inhibits GYS2 expression via its 120-134 aa region.

**Figure 7.**
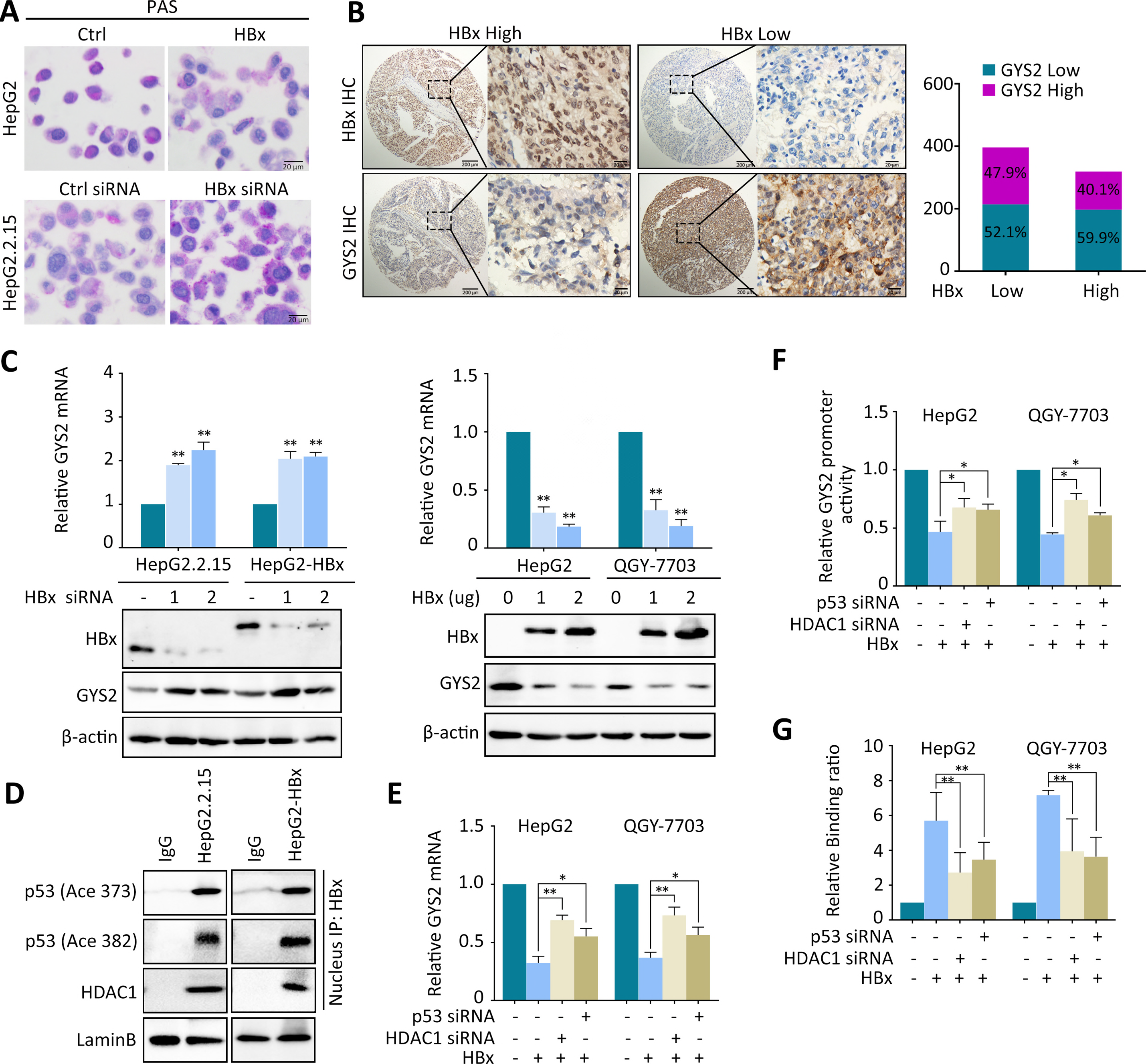
HBx/HDAC1 complex synergistically enhanced the downregulation of GYS2 by p53. **A.** Glycogen content were shown by PAS staining with HBx overexpression in HepG2 and silenced in HepG2.2.15. **B.** HBx and GYS2 expression were analyzed in TMA cohort by IHC. Representative IHC photographs of high and low HBx groups and corresponding GYS2 images were presented. The correlation between HBx and GYS2 were calculated and GYS2 proportion were shown as ratio in high and low HBx groups. **C.** HBx was silenced by siRNAs in HepG2.2.15 and HepG2-HBx cells and overexpressed by pcDNA3.1-HBx-HA plasmid in HepG2 and QGY-7703 cells. GYS2 mRNA and protein expression was examined by qRT-PCR and western blot. **D.** The interaction between HBx and acetylated p53 as well as HDAC1 was detected by co-IP in nucleus. **E.** The expression of GYS2 mRNA were detected by qRT-PCR with the depletion of HDAC1/p53 and overexpression of HBx. **F.G.** GYS2 promoter activity (G) and promoter binding ratio (h) were determined with the depletion of HDAC1/p53 and overexpression of HBx. ^*^*P*<0.05, ^**^*P*<0.01.

It has been reported that HBx’s repressive activity requires the recruitment of collaborative factors, and that histone deacetylases (HDACs) are the major gene suppressors that interact with HBx (Chung & Tsai, 2009; Yoo et al, 2008). We treated HepG2.2.15 and HepG2-HBx cells with siRNAs for HDAC family members. GYS2 mRNA expression was induced only in cells with HDAC1 depletion (Supplementary Fig. 13). Co-IP experiments presented the interactions among HBx, HDAC1 and acetylated p53 in nucleus (Fig. 7D). In cells with HBx overexpression, knockdown of HDAC1 or p53 partially attenuated the reduction of GYS2 mRNA (Fig. 7E). Luciferase and ChIP assays revealed that siRNA for HDAC1 or p53 reduced the binding of HBx to GYS2 promoter (Fig. 7F-G). These data suggest that HBx works with HDAC1 to facilitate the p53-mediated suppression of GYS2 in HBV-positive HCC.

## Discussion

An emerging theme in cancer biology is that aberrant metabolic reprogramming linked to oncogenic transformation or tumor progression (Schulze & Harris, 2012). However, rare attention has been paid to glycogen metabolism. Liver, as the main organ of glycogen deposition, shows dynamic and aberrant glycogen content during carcinogenesis (Toshkov et al, 1994). The regulation of glycogen in liver seems different to other kinds of tissues with unknown mechanisms (Lea et al, 1972; Rousset et al, 1981). Here, we reported for the first time that the glycogen content was dramatically decreased in HCC tissues and correlated with unfavorable patient outcome, which was attributed to the newly identified signaling axis HBx/GYS2/p53. As the rate-limiting enzyme of glycogen biosynthesis primarily expressed in liver, GYS2 represents a promising therapeutic target for the clinical management of HCC.

The clinical significance, biological function, and molecular regulation of GYS2 were investigated in this study. Loss of GYS2 significantly correlated with decreased glycogen content and unfavorable patient outcomes in a large cohort containing 763 patients with HCC. GYS2 silencing promoted HCC cell proliferation but not migration by targeting p53 signaling pathway. Our findings provided two separate lines of evidence supporting the relationship between GYS2 and p53. On one hand, we identified p53 as the downstream target of GYS2. Restoring expression of p53 partly attenuated cell growth and colonies formation promoted by GYS2-depletion. In p53-depleted cell line Hep3B, neither knockdown nor ectopic overexpression of GYS2 influenced the cell viability (Supplementary Fig.14). Mechanically, we found that GYS2 interacted with p53 in the cytoplasm with the competition of MDM2 to stabilize p53 from proteasomal degradation. On the other hand, p53 suppressed GYS2 expression and glycogen content in a negative feedback manner. p53-mediated transcriptional repression is an area much less understood than its transcriptional activation. To the best of our knowledge, p53 is known to suppress gene expression by recruiting repressive complex HDACs or polycomb proteins to specific gene promoters (Lin et al, 2005; Murphy et al, 1999; Zeng et al, 2011). In our studies, the acetylation of p53 at K373 and K382 was catalyzed by p300 and enhanced by the presence of GYS2, which consequently strengthened the DNA-binding ability of p53 to GYS2 promoter, by the supporting of a repressive transcriptional complex with HBx and HDAC1. Taken together, these data demonstrate that p53 and GYS2 may exhibit functions in HCC via equilibrating the expression of each other.

Previous studies have demonstrated the pivotal role of MDM2 in modulating p53 expression. For example, CD147 inhibited p53 at post-transcriptional level through lactate-mediated PI3K/Akt/MDM2 pathway (Huang et al, 2014). PHLDB3 facilitated ubiquitination and degradation of wild-type p53 in dependence on MDM2(Chao et al, 2016). MDM2 is an E3 ubiquitin ligase that directly binding to the N-terminal 1-42 aa of p53 to induce ubiquitin-mediated proteasomal degradation (Inoue et al, 2016; Nomura et al, 2017). Interestingly, our findings showed that GYS2 directly interacted with MDM2 and competitively binding to 1-101 aa region of p53 to prevent MDM2-dependent p53 ubiquitination.

As we know that p53 is frequently mutated in HCC, the effect of GYS2 on mutated p53 should be studied. Here, we found that GYS2-depletion increased p53 protein in Huh7 and PLC/PRF/5 cell lines bearing mutated p53 (Supplementary Fig.15A). *In vitro* assays indicated that GYS2 also influenced the cell viability in these two cell lines (Supplementary Fig.15B-D). Clinical data in TCGA suggested the correlation between GYS2 deficiency and unfavorable overall survival in p53 mutated cohorts (Supplementary Fig.15E). However, the detailed role of GYS2 in p53 mutated cells remain uncovered. Whether GYS2 regulate mutated p53 via MDM2 or other epigenetic ways required further study.

The synthesis and breakdown of glycogen are dependent on specific enzymes and various regulatory proteins. p53 has been reported as the crucial regulator of glucose metabolism by suppressing glucose uptake and glycolysis in tumor cells (Jiang et al, 2011). However, the involvement of p53 in glycogen regulation remains obscure. Here, we identified that p53 overexpression reduced whereas knockdown of p53 induced the accumulation of glycogen in HCC cells. However, this effect was abolished in cells with GYS2 depletion. Besides of GYS2, p53 was capable of transcriptionally regulating other genes involved in glycogen synthesis, such as PYGL, GBE1, G6PC and GSK3β. Hence, these results suggest p53 as a novel mediator of glycogen regulation.

HCC is a chronic inflammation-related cancer that develops primarily from chronic hepatitis B virus infection. Previous studies indicated the involvement of HBV in liver glycogen regulation (Lu et al, 2013; Toshkov et al, 1994). It has been reported that HBx requires the recruitment of collaborative molecules to exert its DNA binding function. We identified HDAC1 as the workmate of HBx for GYS2 repression, in line with previous data that the interaction between HBx and HDAC1 led to the repression of the target genes (Chung & Tsai, 2009; Yoo et al, 2008). Meanwhile, we also found that HBx interacted with acetylated p53 in the nucleus, which enhanced the p53-mediated GYS2 suppression. It is known that HBx modulate the activity of GSK-3β and GSK-3β was also the upstream regulator of GYS2 (Wang et al, 2012; Wang et al, 2015; Yang et al, 2009). However, we found that HBx transcriptionally downregulate GYS2 independent of GSK-3β (Supplementary Fig. 16). As a result, the repressive complex HBx-HDAC1-p53 collaboratively bound to the promoter of GYS2 and inhibited the expression of GYS2, resulting in the decrease of glycogen content.

In summary, our study demonstrate that GYS2 is dramatically downregulated and accompanied by loss of glycogen in HCC. The newly identified signaling axis HBx/GYS2/p53 provided novel mechanistic insight into explaining glycogen metabolic disorders during tumor progression and complemented current understandings of metabolic reprogramming. Our results suggest GYS2 as a feasible prognostic biomarker and therapeutic target for HCC patients.

## Material and Methods

### Patients, tissue specimens and follow-up

A total of 768 primary HCC samples and their corresponding non-tumor tissues were obtained from HCC patients who underwent hepatectomy at Sun Yat-Sen University Cancer Center (SYSUCC). All pathological specimens were collected along with complete clinical and pathological data. Archived paraffin-embedded specimens were selected and re-embedded into new paraffin blocks for tissue microarray (TMA). Another 69 HCC cases with portal vein embolus were recruited between August 2011 and August 2012. This study was approved by the Institute Research Medical Ethics Committee of Sun Yat-Sen University Cancer Center. None of the patients had received radiotherapy or chemotherapy before surgery. All samples were anonymous.

### Histology and glycogen detection

The TMA blocks were cut into 4 μm sections to undergo H&E staining and immunohistochemistry (IHC) staining. Protein expression levels of GYS2 and p53 stained TMA slides were assessed by two independent pathologists (Shi-Xun Lu and Li-Li Liu). Staining intensity multiply by proportion was calculated as the IHC score. Staining intensity was recorded as four grades (0, 1, 2, and 3) and proportion was recorded as five grades (0, 1, 2, 3, and 4). The median of the IHC score was chosen as the cut-off value. For periodic acid-Schiff (PAS) staining, the tissues were treated with periodic acid solution for 5 min and then covered with Schiff’s reagents for 10 min according to the protocol. High PAS group including positive cells >10% or cluster staining while dot or ≤10% staining were assigned as low PAS group. Specimens were pretreated with diastase (D-PAS) as the control of PAS staining. The glycogen content in fresh tissues and cell lines was quantitated by glycogen assay kit ab65620.

### Cell culture

Bel-7402, SMMC-7721, HepG2, QGY-7703, HepG2.2.15 and Hep3B HCC cells as well as immortalized human liver cell line L-02 and QSG-7701 were obtained from the Type Culture Collection Cell Bank, Chinese Academy of Science Committee (Shanghai, China) and routinely cultured in Dulbecco’s modified Eagle’s medium (DMEM) supplemented with 10% fetal bovine serum (Gibco, south American). HepG2-HBx cells stably expressing GFP tagged HBx were established in our lab (Chen et al, 2017a). All cells were maintained in a humidified incubator at 37 °C and 5% CO_2_.

### Inhibitors

Inhibitors Nutlin-3 (HY-50696), CHIR-98014 (HY-13076), CHX (HY-B1248) and MG-132 (HY-13259) were purchased from MedChemExpress Company and added to the culture cells in indicated times and concentrations.

### Western Blot

Total proteins or nucleus and cytoplasm protein were extracted from cells using lysis buffer (Beyotime Biotechnology Shanghai, China) supplemented with protease inhibitor. Western blot was performed with the standard method as previously described (Chen et al, 2017b). Antibodies used in this study were GYS2 (1:500, Sigma, USA), p53 (1:2000, Santa Cruz, CA, USA), HA (1:1000, Santa Cruz, CA, USA), β-actin (1:2000, Santa Cruz, CA, USA), MDM2 (1:2000, Santa Cruz, CA, USA), p300 (1:500, Santa Cruz, CA, USA), 14-3-3σ (1:1000, Santa Cruz, CA, USA), CylinD1 (1:1000, Santa Cruz, CA, USA), p21 (1:2000, CST, USA), CylinA (1:1000, Santa Cruz, CA, USA), CylinE (1:1000, Santa Cruz, CA, USA), HDAC1(1:2000, CST, USA), p53(ace373) (1:1000, Abcam, UK), p53(ace382) (1:1000, Abcam, UK), p53(ace120) (1:500 SAB signalway antibody, USA), p53(ace320) (1:500, SAB signalway antibody, USA)

### Quantitative real-time RT-PCR (qRT-PCR)

Total RNA isolated by Trizol reagent (BIOO Scientific Co., USA) was reverse transcribed using M-MLV Reverse Transcriptase (Promega Inc., USA). SYBR green-based quantitative real-time PCR (Vazyme Biotech, Nangjing, China) was carried subsequently. The sequences of primers shown in Table S1.

### Plasmid construction and RNA interference

Plasmids encoding GYS2 and p53 were cloned into the recombinant plasmids pcDNA 3.1/hygro(+) empty vector. Their function were confirmed by sequencing and their expression level was assayed in HCC cells. The plasmids were transfected into HCC cell lines using Lipofectamine™ 2000 (Invitrogen, Life Technologies, USA) reagent. Small interfering RNAs (siRNAs) targeting GYS2 were purchased from Santa Cruz (SC-69704). Others were designed from Shanghai GenePharma Co. Ltd (Shanghai, China) in Table S1. Transfection was performed by using the Lipofectamine™ RNAiMAX (Invitrogen, Life Technologies, USA).

### Migration assay

Cell motility was assessed by cell migration assay. 2 × 10^4^ cells were plated in the upper compartment of transwell chambers (8-μm pore size, Millepore, Darmstadt, Germany) in serum-free medium. Fresh media containing 10% FBS was placed in the lower chamber. After incubation for 24-48 h, cells on the lower membrane were fixed by 20% methanol and staining with 0.1% crystal violet then counted under a microscope. The experiments were performed in triplicate and repeated three times.

### MTT and colony formation assays

2× 10^3^ cells were seeded in 96-well plates with 100 μl medium and cultured for 5 days. MTT stock solution was added to each well by 10 μl/well for 4 h at 37°C. After addition of DMSO (150 μl/well), absorbance at 490 nm was measured. For the colony formation assay, 500 cells were seeded into 6-well plates and incubated for 14 days. Colonies were fixed with methanol, stained with 0.1% crystal violet and counted.

### Animal model

Four-weak-old male BALB/c nude mice were purchased from Vital River Company (Beijing, China). Mice were randomized into each group and subcutaneously inoculated with HCC cells stably transfected with GYS2 shRNA or overexpression plasmids. Tumor growth was monitored every three days. Mice were sacrificed 27day after inoculation. Tumor volume was calculated with this formula: tumour volume (mm^3^) X (length_width^2^)/2. Tumor were fixed into paraffin-embedded specimens to detect glycogen content and proliferation rate by PAS staining and Ki-67 staining. All animal studies were approved and performed by the animal institute of Sun Yat-sen University Cancer Center according to the protocols approved by the Medical Experimental Animal Care Commission of Sun Yat-sen University Cancer Center.

### EdU assay and Immunoflorescence staining

For EdU assay, cells were pre-cultured with EdU for 3 hours using a mixture reagent Kit (Keygene Biotech, Jiangsu, China) following the mafufacture structure. For immunoflorescence staining, cells were washed twice in PBS, then fixed in 3.7% formaldehyde and permeabilized with 0.1% Triton X-100 both for 10 min at room temperature. After blocked in 1% bovine serum albumin in PBS for 30 min, incubated the cells with diluted primary antibody overnight. After washing with PBS three times, secondary antibody were added for 1hour incubation at room temperature for. DAPI solution were applied and imaged were captured using confocal microscope.

### Flow cytometry assay

Cells transfected with siRNAs or plasmids were washed with flow buffer and stained with respective dye (Annexin V and propidium iodide) in the dark according to the manufacture (11988549001, Roche). The cells were then analyzed using Beckmanculter flow cytoflex and Modfit (Verity, Topsham, ME, USA) software programs.

### Immunoprecipitation (Co-IP) and Chromosome immunoprecipitation (ChIP)

For Co-IP assay, protein were harvested in lysis buffer (Beyotime Biotechnology Shanghai, China) supplemented with protease inhibitor (P-8340, Sigma, USA). After culturing with primary antibody as indicated in the figure legends or mouse IgG and for 4h, protein A/G PLUS beads (sc-2003, Santa Cruz, CA, USA) were added and incubated overnight. Precipitation were washed at least three times with lysis buffer. For ChIP assay, all the procedures were conducted following the guideline of manufacture (Pierce™Magnetic ChIP Kit 26157). The primers used in ChIP assay were described in Table S1.

### Dual-luciferase reporter assay

For the dual-luciferase report assay, HepG2 cells were transfected with GYS2 promoter or empty vector in pGL3-basic separately, each co-transfected with pcDNA 3.1-HA-HBx plasmid. Renilla luciferase activity was employed as an internal control. Luciferase activity was analyzed with the Dual-luciferase Reporter Assay System (Promega, CA, USA).

### Statistical analysis

Data are shown as means ± standard deviation. Statistical analyses were performed with the SPSS 19.0 software (SPSS, Chicago, IL, USA). Student’s t-test, Pearson’s χ2 test, Fisher’s exact test, Kaplan-Meier method and Multivariate Cox proportional hazards regression model were conducted accordingly. *P* < 0.05 (two-tailed) was considered statistically significant.

## The paper explained

### Problem

Hepatocellular carcinoma (HCC) is a leading cause of cancer death globally. Emerging evidence has strongly suggested that metabolic reprogram plays a critical role in HCC progression. However, abnormalities in glycogen deregulation in HCC are rarely studied. The clinical significance, biological function, as well as molecular regulation of glycogen and liver glycogen biosynthesis enzyme GYS2 were investigated in this study.

### Results

Our preliminary data of high through-put sequencing revealed that the expression of GYS2 was dramatically downregulated in HCC tissues, which correlated with decreased glycogen content and nfavorable patient outcomes. GYS2 overexpression suppressed, whereas GYS2 knockdown facilitated cell proliferation *in vitro* and tumor growth *in vivo* in a p53-dependent manner. GYS2 prevent p53 from MDM2-mediated ubiquitination and enhanced p300-induced acetylation of p53 at K373/382, which in turn inhibited the transcription of GYS2 in the support of HBx/HDAC1 complex. HBx/p53 collectively restrict the accumulation of glycogen in HCC in the presence of GYS2.

### Impact

Collectively, our findings suggest GYS2 as a prognostic factor and functions as a tumor suppressor in HCC. The newly identified HBx/GYS2/p53 axis is responsible for the deregulation of glycogen metabolism and represents a promising therapeutic target for the diagnostic and treatment of HCC.

## Acknowledgements

This work was supported by grants from the National Key R&D Program of China (2017YFC1309000), The National Natural Science Foundation of China (No. 81572405, 81572406, 81502079, 81602135), The National Natural Science Foundation of Guangdong province (No.2018B030311005) and Science and technology program of Guangzhou (201707020038).

## Author contributions

Conception and design of the study: Shi-Lu Chen, Chris Zhiyi Zhang, Jing-Ping Yun; Generation, collection, assembly, analysis of data: Shi-Lu Chen, Ying-hua Pan, Chun-Hua Wang, Yang-fan He, Cen-Shan Lin; Scoring and evaluation of IHC stained slides: Li-Li Liu, Shi-Xun Lu, Xia Yang; Drafting and revision of the manuscript: Shi-Lu Chen, Chris Zhiyi Zhang, Dan Xie, Jing-Ping Yun; Approval of the final version of the manuscript: all authors.

